# *Pseudomonas putida* group species serve as reservoirs of conjugative plasmids disseminating Tn*402*-like class 1 integrons carrying *bla*_VIM-2_ metallo-β-lactamase genes

**DOI:** 10.1101/2020.12.23.424275

**Authors:** Marco A. Brovedan, Patricia M. Marchiaro, María S. Díaz, Diego Faccone, Alejandra Corso, Fernando Pasteran, Alejandro M. Viale, Adriana S. Limansky

## Abstract

The *Pseudomonas putida* group (*P. putida* G) is composed of at least 21 species associated to a wide range of environments, including the clinical setting. Here, we characterized 13 carbapenem-resistant *P. putida* G clinical isolates carrying *bla*_VIM-2_ from different hospitals of Argentina. Multilocus sequencing (MLSA) and phylogenetic analyses based on the 16S rDNA, *gyrB* and *rpoD* sequences comparison allowed us to assign them to 7 well-differentiated species. Sequencing analysis revealed that *bla*_VIM-2_ genes were carried in these isolates by three different class 1 integrons (In*41*, In*899* and In*528*) embedded into Tn*402*-like transposons. Those harboring In*41* and In*899* were designated Tn*6335* and Tn*6336*, respectively, with the former found among 10 isolates. Both encompassed complete transposition modules and inverted repeats boundaries characteristic of the Tn*5053*/Tn*402* family, whereas the third, bearing In*528*, exhibited a defective *tni* module. Tn*6335* and Tn*6336* were located in conjugative pLD209-type plasmids in *P. asiatica, P. juntendi, P. putida* G/II, and *P. putida* G/V isolates, and could be mobilized to *Escherichia coli* and *P. aeruginosa* indicating a relevant mechanism of *bla*_VIM-2_ dissemination. In other *P. asiatica* and *P. putida* G/II isolates, Tn*6335* was found inserted into the Tn*21* subgroup transposons-*res* region, indicating capability for intragenomic mobilization and further dissemination associated to Tn*3* family transposons. The Tn*402*-like defective element was also found inserted into the *res* region of another Tn*3* family transposon in a *P. monteilii* isolate, but in an atypical orientation. Overall findings shed light on the mechanisms by which resistance genes move through environmental and opportunist *Pseudomonas* species.

## INTRODUCTION

The members of the genus *Pseudomonas* are ubiquitous in nature, and thrive in very different ecological niches including soils, water, sediments, air, and human environments (1, 2, 3, 4, 5). *Pseudomonas* members are endowed with a wide metabolic versatility and a broad potential for adaptation to challenging environmental conditions. The genus includes pathogenic species such as *P. aeruginosa*, an important cause of healthcare associated infections especially affecting immunocompromised patients (6). One main cause of the success of *P. aeruginosa* in the nosocomial environment is represented by its ability to resist the most classes of antimicrobial agents of clinical use (multidrug resistance, MDR), a situation that nowadays includes last-resource therapeutic options such as the carbapenems. Horizontal gene transfer (HGT) of genetic platforms carrying resistance genes from other bacteria inhabiting the same ecological niche has certainly impacted the *P. aeruginosa* ability to evolve MDR and to persist in human-associated environments (6).

Different phenotypic and chemotaxonomic features have been extensively used in *Pseudomonas* classification, but much more reliable approaches are represented by genomic-based procedures (1, 2, 3, 4; 5). In this context, while 16S rDNA sequence comparisons have been pivotal to delineate the limits of the *Pseudomonas* genus, the discriminatory ability of this approach at the intrageneric level is generally low (3). A more accurate definition at the species level requires other approaches such as multilocus sequence analysis (MLSA) using a number of selected core genes that include, besides 16S rDNA, other housekeeping genes such as *gyrB, rpoB* and/or *rpoD* (1, 2, 3, 4, 5). This is the case of the *P. putida* group (*P. putida* G), which is composed of at least 21 assigned species which share with *P. aeruginosa* a wide range of environmental niches including soils, freshwater, and animals (1, 2, 3, 4, 5, 7, 8, 9; 10). Among this group, *P. putida, P. monteilii, P. fulva, P. mosselii*, and newly proposed species such as *P. asiatica* and *P. juntendi* have been recently associated to infections affecting human hosts (4, 5, 11, 12, 13, 14). Although most *P. putida* G isolates of human origin have generally exhibited susceptibility to most clinically-employed antimicrobial drugs, an emerging resistance to carbapenems among them has more recently been reported. This, in turn, has been accompanied by the detection of metallo-β-lactamase (MβL) genes of the VIM, IMP, DIM, or NDM families, pointing to *P. putida* G members as possibly serving as active reservoirs of such resistance genes (13, 15, 16, 17, 18, 19, 20, 21, 22, 23, 24, 25). The potential dissemination of these genes to co-existing human pathogens including *P. aeruginosa* and members of the *Enterobacteriaceae* family poses a serious challenge to successful antimicrobial therapy (6, 13, 17, 26).

Among MβL genes, *bla*_VIM-2_ is one of the most widely found among clinical strains of *P. aeruginosa* worldwide (26, 27, 28). In MDR *P. aeruginosa* strains the *bla*_VIM-2_ gene is usually carried by “mobile” class 1 Tn*402*-like integron, which are members of the Tn*5053*/Tn*402* family (29, 30, 31, 32, 33). The members of this transposon family are known by characteristic 25-bp initial and terminal inverted repeats (IRi and IRt, respectively) boundaries, a *tni* module composed of *tniA*, *tniB*, *tniQ*, and *tniC* (also designated *tniR*) genes responsible for replicative transposition, and to generate a 5-bp direct repeat (DR) at the site of insertion. Moreover, they show a notable selectivity for recombination (*res*) regions located upstream of *tnpR* genes of Tn*3* family members such as Tn*21*, or to the equivalent regions associated to segregational mechanisms of particular plasmids (32, 33, 34, 35, 36, 37). In *P. aeruginosa*, *bla*_VIM-2_-containing Tn*402* transposons are located in the chromosome or, more worryingly, associated to conjugative plasmids which may enormously facilitate the spread of carbapenem resistance (6, 13, 18, 31). In contrast to the case of *P. aeruginosa*, however, fewer data exist at present on the genetic platforms carrying *bla*_VIM-2_ genes among other *Pseudomonas* species sharing similar niches, including *P. putida* G. A detailed characterization of these platforms could then provide important insights into the role of *P. putida* G members as reservoirs of *bla*_VIM-2_ genes, and on the mechanisms of spreading of carbapenem and other resistance genes among clinically relevant bacterial species.

We have recently characterized by complete sequencing a self-transferable plasmid, pLD209, housed by the carbapenem-resistant *P. putida* G clinical strain LD209 (13, 17). pLD209 carries a Tn*402* integron/transposon possessing *bla*_VIM-2_ and *aacA4* gene cassettes (13, 17). In this work, we characterized different genetic platforms carrying *bla*_VIM-2_ genes present in a collection of carbapenem-resistant *P. putida* G clinical strains isolated in hospitals of two major cities of Argentina. The observations reported here led us to propose intra- and inter-genomic mechanisms of *bla*_VIM-2_ dissemination among environmental *Pseudomonas* and Gram-negative pathogens co-existing in the clinical setting.

## RESULTS

### Bacterial isolates

A total of 13 carbapenem-resistant clinical isolates from hospitals of the Buenos Aires City and Rosario City areas of Argentina, identified phenotypically as belonging to the *P. putida* group by the VITEK 2C System, were included in this study (Table 1 and Table S1). All of the *P. putida* G isolates analyzed here showed clinical resistance to imipenem and meropenem, and to other β-lactams such as ceftazidime and piperacillin-tazobactam. With the exception of BA9115, all other isolates showed resistance to gentamicin. Ten isolates were also resistant to ciprofloxacin (Table S1). Concerning carbapenem resistance, an EDTA-imipenem microbiological assay in combination with an EDTA disk synergy test (38) revealed the presence of MβL activity in all 13 isolates. The searching of different MβL genes by PCR employing specific primers for *bla*_IMP_, *bla*_VIM_, *bla*_SPM_ *and bla*_NDM_ genes (Table S2), followed by sequencing analysis of the obtained amplicons, found only *bla*_VIM-2_ in all 13 isolates.

**TABLE 1.**
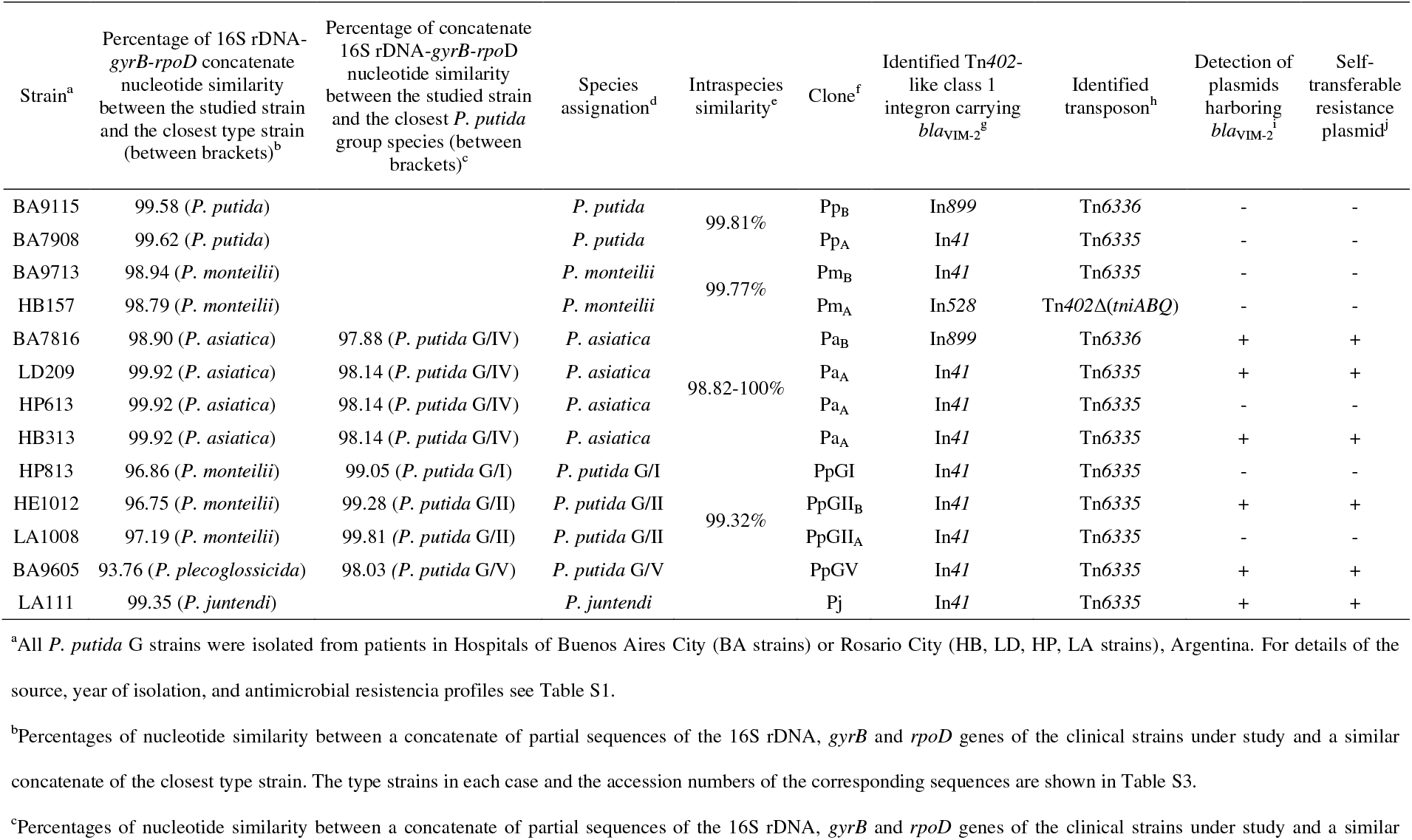
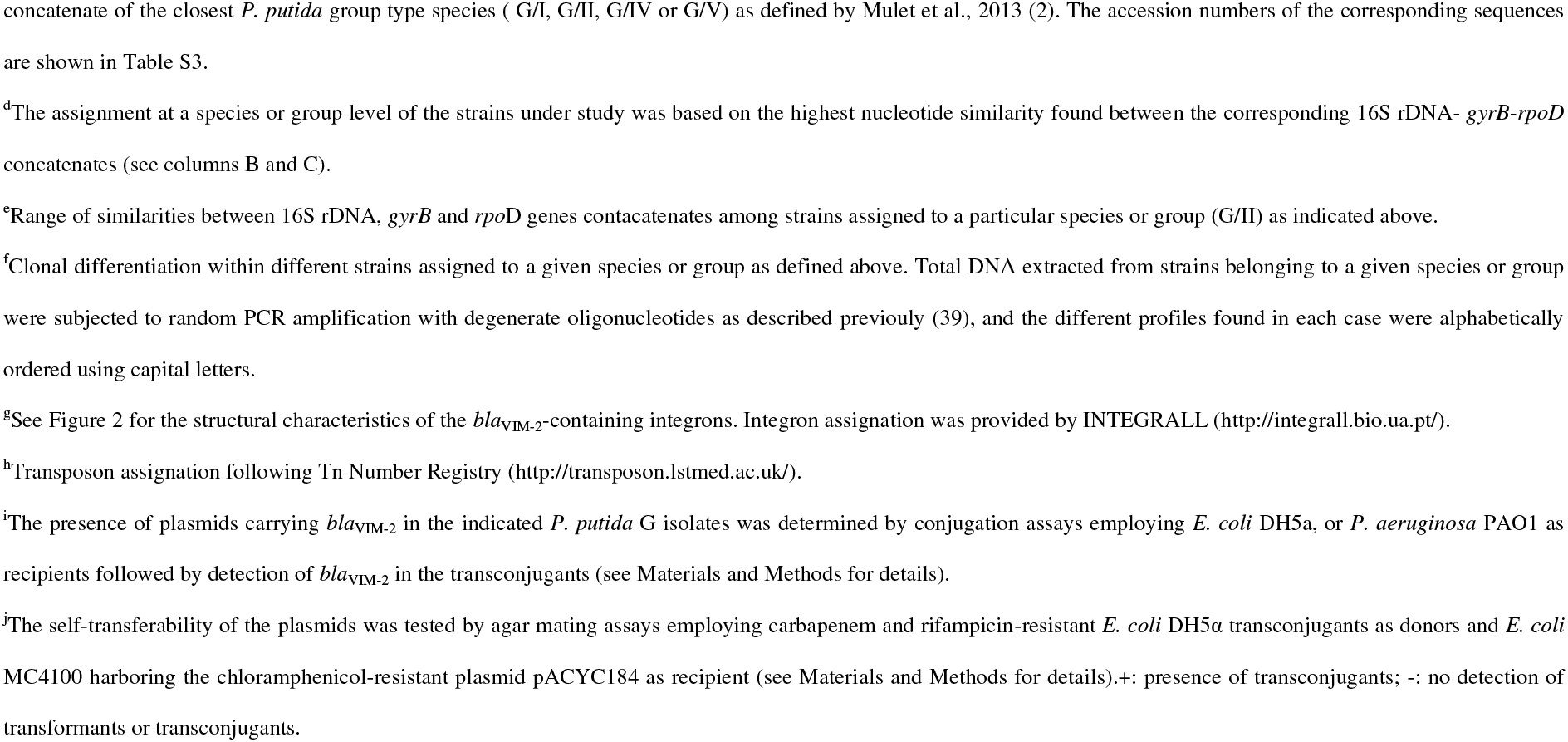
Strains of *P. putida* group used in this study

### Assignment of the analyzed isolates to the species level within the *P. putida* G

A more precise assignment of the 13 isolates was conducted by MLSA (2) employing comparisons of the concatenated partial sequences of 16S rDNA (1,301 bp), *gyrB* (669 bp) and *rpoD* (677 bp) genes. After obtaining the corresponding amplified sequences for each isolate (Table S2), the derived concatenates were first aligned and compared to the equivalent sequences extracted from reference strains representing the different species of the *P. putida* G, including recently proposed species (1, 2, 4, 5). The threshold percentage of identity between concatenate sequences used to discriminate between species within *P. putida* G was set at ≥ 97.5 % (1, 2). In this study, we assigned each of the isolates described here to the species to which the corresponding concatenates shared the higher percentage of sequence identity (see summary in Table 1). Thus, isolates BA7908 and BA9115 were assigned to *P. putida sensu stricto;* HB157 and BA9713 to *P. monteilii*; BA7816, LD209, HB313 and HP613 to *P. asiatica*; and LA111 to *P. juntendi*, as judged by percentages of identity that ranged from 99.3% (LA111, *P. juntendi*) to 99.94% (BA9713, *P. monteilii*) to the corresponding species (Table 1). In the case of 4 isolates (HP813, HE1012, LA1008, and BA9605) the highest percentages of concatenate sequence identity to the corresponding closest defined species were found to be lower than the 97.5% threshold value (Table 1). However, as also seen in the Table, these isolates could be assigned to recently proposed *P. putida* G novel species (1) as judged by percentages of concatenate sequence identity that ranged from 98.03 to 99.81%. This included the assignment of LA1008 and HE1012 to *P. putida* G/II, HP813 to *P. putida* G/I, and BA9605 to *P. putida* G/V (Table 1). Also, intraspecies sequence similarity values of different isolates assigned to the same species (*P. putida*, *P. monteilii*, and *P. putida* G/II, Table 1) were found to be higher than 98.8%.

Secondly, a maximum-likelihood (ML) phylogenetic analysis using the corresponding concatenated sequences of 16S rDNA, *gyrB* and *rpoD* genes validated the species assignations obtained above (Fig. 1). This analysis also included the corresponding concatenates from additional type strains of species of the *P. putida* G (1, 2, 4, 5). *P. aeruginosa* (ATCC 10145) and *P. oryzihabitans* (ATCC 43272) were used as outgroups (Fig. 1, see Table S3 for details). This analysis confirmed the assignation of isolates BA9115 and BA7908 to *P. putida*; HB157 and BA9713 to *P. monteilii*; BA7816, LD209, HP613, and HB313 (the latter three isolates displaying identical concatenate sequences, Table S3) to *P. asiatica*; and LA111 to *P. juntendi*, as judged by the monophyletic groups formed in each case with the corresponding defined species. The clinical strain LD209, previously characterized as belonging to the *P. putida* G on the basis of phenotypic procedures (13), could now be more confidentially assigned to *P. asiatica* on the basis of the analysis described above.

**FIG 1.**
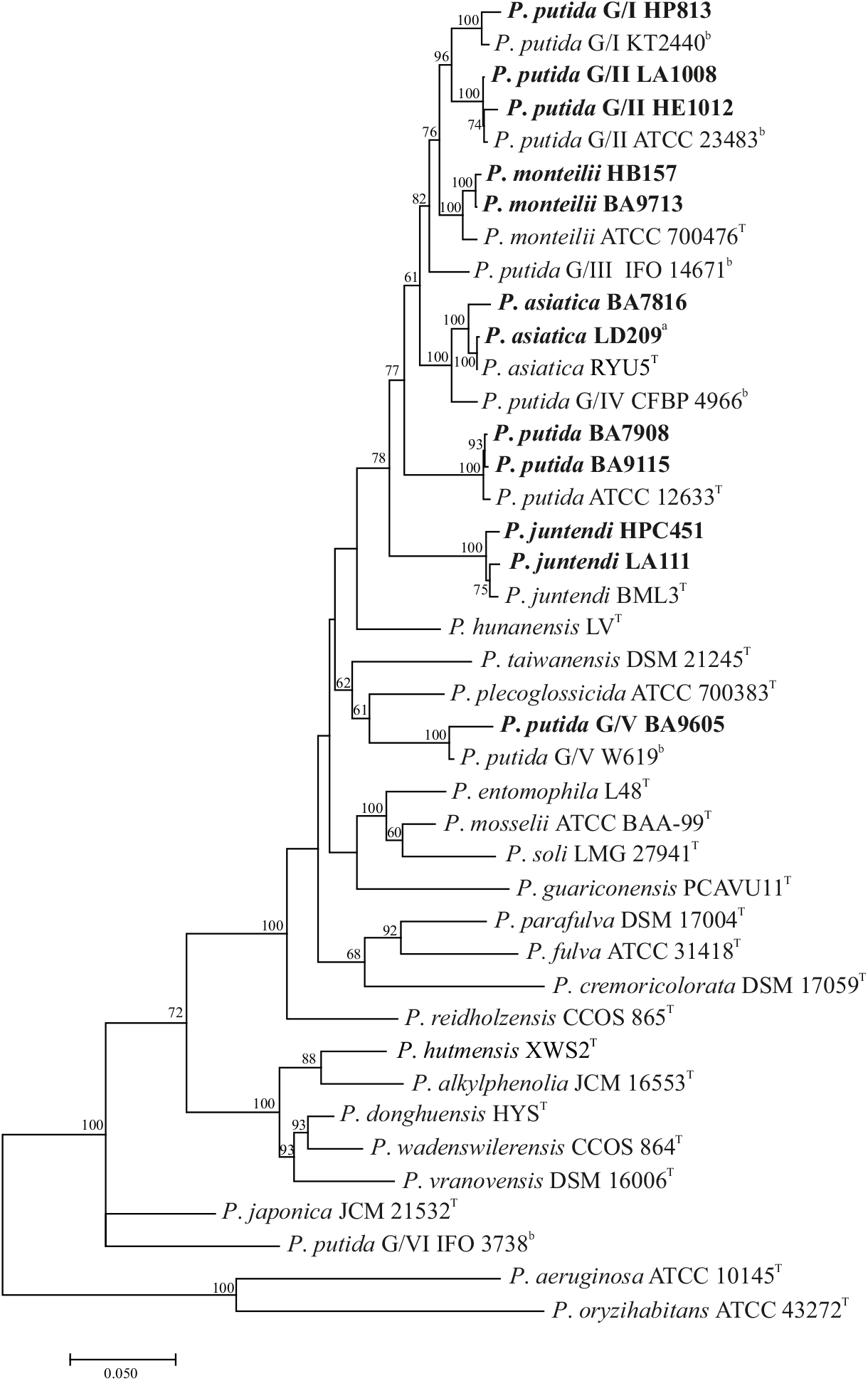
Phylogenetic analysis of the *Pseudomonas putida* G isolates analyzed in this work. A ML phylogenetic tree was constructed from alignments of the concatenated sequences of defined partial regions of 16S rDNA, and *gyrB* and *rpoD* genes corresponding to the different *Pseudomonas* spp. strains indicated in the figure. The *P. putida* G clinical isolates analyzed in this work are in bold. The analysis also incorporates the corresponding concatenated sequences of 21 *P. putida* G type strains which have received species assignation (2, 4, 5) as well as from 6 strains representing proposed novel species (*P. putida* G/I to *P. putida* G/VI, 2, indicated by ^b^ superscripts). In addition, the concatenate sequences of *P. aeruginosa* ATCC 10145^T^ and of *P. orzihabitans* ATCC 43272^T^ type strains were included to serve as outgroup sequences of the *P. putida* group. For details on the corresponding nucleotide accession numbers of the sequences employed here see Table S3. Only bootstrap percentages higher than 60 % (1,000 replicates) are indicated at the nodes. In LD209^a^, the superscript indicates the representative isolate of the clonal lineage Pa_A_ among the *P. asiatica* isolates, which also includes HP613 and HB313 (not shown in the figure). These three isolates share identical sequences of core gene concatenate sequences. The superscript ^T^ indicate the type strains. See Materials and Methods for details.

Of note, some isolates clustered with newly proposed species, such as HP813 with *P. putida* G/I KT2440; LA1008 and HE1012 with *P. putida* G/II ATCC 23483; and BA9605 with *P. putida* G/V W619 (Fig. 1). Thus, our local collection of clinical isolates was composed of (at least) seven different species now assigned to the *P. putida* group. It is worth noting that in the ML tree (Fig. 1) the RYU5 strain, recently proposed as representative of the new species *P. asiatica* within the *P. putida* group (4), clustered with the CFBP4966 strain. The latter was in turn proposed as a representative of a new *P. putida* group species designated G/IV (1). Our observations, including the 98.1% nucleotide identity shown by the corresponding concatenate sequences (Table 1), strongly point to these two strains as belonging to a same species.

Further discriminatory fingerprinting analyses (39) showed that the four clinical isolates assigned here to *P. asiatica* (see above) could be subdivided into two separate clonal lineages: one constituted by LD209, HB313, and HP613, and the other by BA7816 (Fig. S1, Table 1). Also, the two isolates assigned to *P. putida sensu stricto* (BA9115 and BA7908, respectively) could also be ascribed to two separate clonal lineages, and the same situation was found for the two isolates assigned to *P. monteilii* (HB157 and BA9713) and the two assigned to *P. putida* G/II (LA1008 and HE1012) (Fig. S1, Table 1). These results indicated the presence, among the 13 *P. putida* G isolates included in this study, of 11 still distinguishable isolates by using more discriminatory genetic procedures (Fig. S1). This not only exemplifies the difficulties in defining the limits between species in this phylogenetically closely-related group (1, 2, 3, 4, 5), but also indicates that further delimitations of the presently-accepted species may be necessary in the future.

In summary, the above results indicate the existence of (at least) seven distinguishable species of the *P. putida* G carrying *bla*_VIM-2_ among our isolates, and reinforce the notion of this group acting as a carbapenem resistance genes reservoir (13, 17). Moreover, they also revealed the persistence and/or dissemination in our local nosocomial environment of a specific clone of *P. asiatica* (designated Pa_A_, Table 1 and Table S1). Similar results concerning the nosocomial transmission of a single clone of *P. putida* have been reported in a tertiary hospital of Pittsburgh, Pennsylvania, U.S.A. (40). A further characterization of the *bla*_VIM-2_-carrying genetic platforms in the above-mentioned isolates could then shed light on the mechanisms involved in their exchange among *P. putida* G species.

### Characterization of genetic structures harboring *bla*_VIM-2_ genes in the *P. putida* G isolates analyzed

The characterization of the near genomic context of *bla*_VIM-2_ using PCR primers designed to hybridize in the 5’ and 3’ conserved segments of “typical” class 1 integrons (Table S2) systematically failed to produce amplification bands in all of the 13 *P. putida* G clinical isolates analyzed here. Conversely, the use of Int1-F and TniC-R2 primers, hybridizing into the 5’-CS and the 5’ region of the *tniC* gene located in the *tni* module of Tn*402* transposons, respectively (Table S2) generated amplification bands in all of them. These results indicated the presence of Tn*402*-like class 1 integrons carrying *bla*_VIM-2_ in all 13 *P. putida* G isolates. Subsequent sequencing analysis of the amplicons not only confirmed the presence of *bla*_VIM-2_-containing gene cassettes and their association to a *tniC* gene of a Tn*5053*/Tn*402* transposon family in all cases, but also identified the presence of three different *bla*_VIM-2_-containing unusual class 1 integrons among them (see below, Fig. 2).

**FIG 2.**
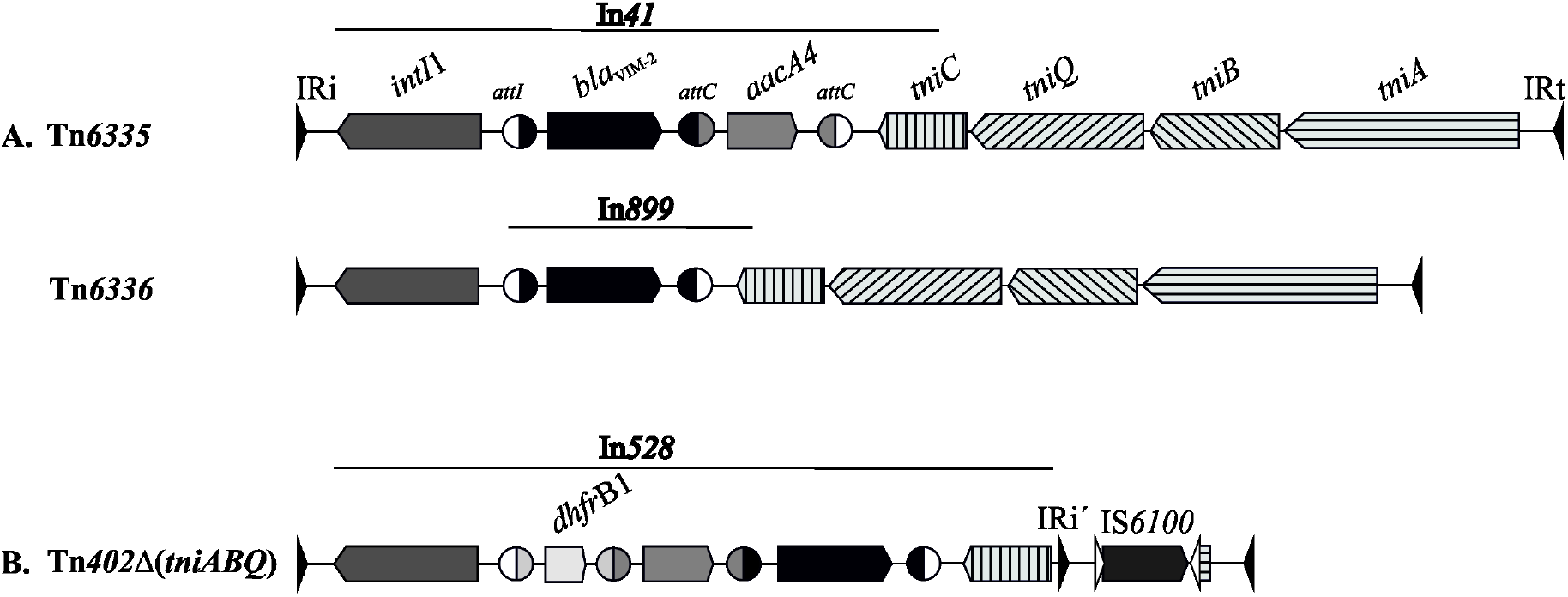
Genetic organization of *bla*_VIM-2_-containing Tn*402*-like class 1 integrons in the *P. putida* G clinical isolates analyzed. **A**. Schematic structure of the class 1 integrons In*41* and In*899* embedded into complete Tn*402*-like transposons designated Tn*6335* (7,633 bp, GenBank accession number GQ857074) and Tn*6336* (6,994 bp, GenBank accession number MN240297.1), respectively. **B**. Same, for the In*528* class 1 integron embedded into an incomplete Tn*402*-like integron detected in *P. monteilii* HB157 and lacking most of the *tni* module (Tn*402*Δ(*tniABQ*), 5,239 bp; GenBank accession number MT192132). The initial inverted repeats (IRi) and terminal inverted repeats (IRt) associated to the left and right boundaries, respectively, of each transposable element are indicated by oppositely-oriented closed arrows at the corresponding Tn borders. The individual genes are represented by boxed arrows (distinctively labeled in each case) that also indicate the corresponding directions of transcription. *IntI1*, integrase (black and gray mosaics); *bla*_VIM_-2, VIM-2 MβL (black); *aacA4*, aminoglycoside acetyl transferase (gray); *tniC*, resolvase (light gray with black vertical stripes); *tniQ/tniB*, auxiliary transposition genes (light gray with black diagonal stripes); *tniA*, transposase (light gray with black horizontal stripes); *dhfrB1*, dihydrofolate reductase (light gray); *IS6100* transposase (dark gray). In the cassette genes located within the integrons, the recombination sites are indicated by two halves of a circle located upstream of and downstream of the associated gene. The two open halves are from the original *attI* site, and halves exhibiting the same shade of gray are from the corresponding *attC* sequences originally associated to a distinctive resistance cassette. Regions on the transposons described in this work sharing significant sequence identity with described mobile elements (see main text for details) are indicated by lines above the corresponding structures. In **B**, the initial inverted repeat (IRi) and terminal inverted repeat (IRt) are indicated by black triangles facing inwards. A second 25-bp sequence identical to IRi, designated IRi’, was also located immediately upstream of the *tniC* gene (black triangle facing IS*6100*). The left inverted repeat (IRl) and right inverted repeat (IRr) of IS*6100* are indicated by the oppositely-oriented open arrows located at the borders of the transposase gene (in dark gray). The structures of Tn*6335* and Tn*6336* were determined by PCR using different pairs of primers (Table S2), followed by sequencing analysis of the amplicons and assembly of the overlapping segments, and further confirmed by complete plasmid sequencing (Fig. 3). The complete sequence of Tn*402*Δ(*tniABQ*) was determined using an inverse PCR procedure. The figure is not drawn to scale. For details see Materials and Methods.

We have previously reported that the *P. asiatica* LD209 strain carries a conjugative plasmid of 38,403 bp, designated pLD209 (13, 17). The detailed analysis of the pLD209 sequence (13) indicated that it harbours a Tn*402*-like transposon of 7,633 bp endowed with a complete *tni* module, and a class 1 integron (In*41*) possessing *bla*_VIM-2_ and *aacA4* resistance cassettes (Fig. 2A). This Tn*402* element has been assigned the denomination Tn*6335* by the Tn Number Registry (41). The sequence characterization of the amplicons obtained from the different *P. putida* G isolates described above indicated the presence of the In*41* arrangement in *P. putida* BA7908, *P. monteilii* BA7913, *P. asiatica* HP613, *P. asiatica* HB313, *P. juntendi* LA111, *P. putida* G/I HP813, *P. putida* G/II HE1012, *P. putida* G/II LA1008, and *P. putida* G/V BA9605 (Table 1). Further PCR amplification of the immediate genetic context of In*41* (see Materials and Methods for details) followed by sequencing analysis confirmed the presence of Tn*6335* in all of these isolates.

Concerning the other isolates, similar procedures than those described above detected only the *bla*_VIM-2_ gene cassette into the variable region of the class 1 integron carried by *P. putida* BA9115 and *P. asiatica* BA7816 (Table 1). This single-cassette integron was previously reported with the designation In*899* (GenBank accession number KJ668595) in *Pseudomonas chlororaphis* M11740 isolated in Argentina, where it was found in a 55-kbp plasmid (42). Further sequencing analysis indicated that In*899* was embedded in a complete Tn*402* transposon of 6,994 bp in our isolates (Fig. 2A). This novel Tn*402* element was assigned the denomination Tn*6336* by the Tn Number Registry (Fig. 2A).

Finally, a similar analysis conducted in *P. monteilii* HB157 (Table 1) indicated a class 1 integron carrying *dhfr*B1, *aacA4* and *bla*_VIM-2_ gene cassettes, accompanied by a defective *tni* module containing only the *tniC* gene (Fig. 2B). Further completion of its downstream region by an inverse PCR procedure (see Material and Methods for details) indicated that it formed part of a Tn*402*-like element spanning a total of 5,239 bp (MT192132.1, Fig. 2B). This methodology allowed to confirm a defective *tni* module composed only of a complete resolvase *tniC* gene and an upstream 44 bp remnant of the intergenic *tniC*/*tniQ* region, accompanied by a short fragment (11 bp) of the 5’ coding region of *tniA* gene (Fig. 2B). The finding of an IS*6100* element near this *tniA* remnant suggests that disruption occurred as the consequence of this IS targeting *tniA*, followed by further deletions/rearrangements at the vicinity of the insertion site. In support to this inference, a BLASTn search indicated that this defective Tn*402*-like element shares almost full identity in the first 4,084 nucleotides (spanning from the IRi at the left boundary to the short 44-bp intergenic fragment located at the 5’ region of the *tniC* gene, and including the integron-associated *dhfrB1-, aacA4-* and *bla*_VIM-2_ gene cassettes) with the equivalent region of a complete Tn*402* family transposon (8,041 bp) located in different *P. aeruginosa* strains including R22 (AM993098.1) and DZ-B1 (KY579949.1). A further stretch of homology spanning 150 nucleotides (81% nucleotide identity) between these elements was found at the corresponding right boundaries, and included the 25-bp inverted repeat IRt and associated sequences (Fig. 2B). Most intriguingly, a second 25-bp inverted repeat identical to IRi was located upstream of the *tniC* gene (designated IRi’ in Fig. 2B). Our database searching identified an identical Tn*402*Δ(*tniABQ*) element in the chromosome of *P. putida* PP112420, a clinical strain isolated in China (GenBank accession CP017073.1; nucleotide positions 5,137,406 to 5,142,644). The observation that this Tn*402*Δ(*tniABQi*) element is bounded by both IRi and IRt and immediate associated sequences characteristic of Tn*5053*/Tn*402* transposons (30) opens the possibility that it could be mobilized to other genomic locations in cells providing in *trans* the enzymes required for transposition. Tn*5053*/Tn*402* family transposons use a replicative mechanism of transposition, in which resolution of cointegrates is mediated by the product of the *tniC* gene acting at a *res* site located in the 61-bp *tniC/tniQ* intergenic region (30). In this context, comparative sequence analysis of the remnant 44 bp region upstream of *tniC* in Tn*402*Δ(*tniABQ*) indicated that the recombination site and immediate adjunt sequences were preserved in this defective element.

### Plasmid transfer of *bla*_VIM-2_-containing Tn*402*-like class 1 integrons from *P. putida* G members to *E. coli* and *P. aeruginosa*

We next analyzed whether the Tn*402* elements characterized above were carried by plasmids endowed with transfer potentiality. For this purpose, conjugation experiments were conducted using the corresponding *P. putida* G isolates as donors, and *E. coli* DH5α or *P. aeruginosa* PAO1 as recipient cells (see Materials and Methods for details). We previously reported that, in laboratory conjugation assays, plasmid pLD209 harbored by *P. asiatica* LD209 could be transferred to *E. coli* DH5α and to *P. aeruginosa* PAO1 (17). By conducting similar conjugation experiments, we could detect plasmid transfer to *E. coli* DH5α and also to *P. aeruginosa* PAO1 when the donor was *P. asiatica* LD209 (used as control), *P. asiatica* BA7816, *P. asiatica* HB313, *P. putida* G/II HE1012, *P. putida* G/V BA9605, or *P. juntendi* LA111, as judged by the selection of imipenem-resistant transconjugants of the corresponding recipients in all of the above cases and the subsequent detection of acquired MβL activity in all of them (Table S4 and data not shown).

The antimicrobial susceptibility patterns of the corresponding *E. coli* DH5α transconjugants (designated Ect209, Ect7816, Ect313, Ect1012, Ect9605 and Ect111, respectively) are shown in Table S4. All of them acquired resistance to imipenem and the other β-lactams tested, as judged by the increments observed in the corresponding MIC values when compared to *E. coli* DH5α cells (Table S4). It is worth noting that, with the exception of Ect7816, the MIC values to gentamicin also increased from 4- to 8-fold in all transconjugants (Table S4). The overall observations are in agreement with the conjugative transfer, from the indicated *P. putida* G isolates, of a plasmid containing Tn*6335* carrying *bla*_VIM-2_- and *aacA4*-resistance determinants in each Ect209, Ect313, Ect1012, Ect9605 and Ect111, and of a plasmid housing Tn*6336* carrying only a *bla*_VIM-2_ cassette in Ect7816 (Fig. 2A).

To further examine the self-transfer capability of the plasmids selected in each of these Ect transconjugants, agar mating assays were done using each of them as donors, and chloramphenicol-resistant *E. coli* MC4100 cells as recipients following previously described procedures (17) (Table 1). The overall results confirmed that the genetic elements under scrutiny (*i. e*., the plasmids derived from Ect313, Ect1012, Ect9605, Ect111 and Ect7816) constitute self-mobilizable plasmids, similarly to the case of pLD209 (17). Thus, all them are seemingly capable of spreading the *bla*_VIM-2_ containing-Tn*402*-like integrons they carry among a wide range of bacterial species.

Plasmids present in the above Ect transconjugants were extracted and subjected to restriction mapping using EcoRI. With the exception of Ect7816, the plasmids derived from Ect1012, Ect9605, Ect313, and Ect111 showed very similar restriction profiles between them, which were in turn very similar to those of Ect209 (data not shown, see also below). Moreover, the obtained sizes corresponded closely to the EcoRI fragment sizes predicted *in silico* from the pLD209 complete DNA sequence (Fig. 3e; 13). The plasmids purified from Ect7816 (henceforth, pBA7816) and Ect111 (henceforth pLA111) were subjected to further sequencing (see Materials and Methods for details) for a subsequent comparative analysis of the obtained structures. pLA111 (GenBank accession number MT192131) was found to be identical to pLD209 (Fig. 3d and 3e, respectively), in agreement with the restriction analysis mentioned above. In turn, pBA7816 (Fig. 3c, GenBank accession number MN240297) was found to be very almost identical (99%) to pLD209, including the replication, transfer, and stability modules. The only main difference between these two plasmids was found in the adaptive module, and consists in the absence of the *aacA4* aminoglycoside resistance cassette in the Tn*402*-like element (Fig. 2A and Fig. 3). It is worth noting that Tn*6336* and Tn*6335* are positioned in equivalent positions in the plasmids, that they are bordered by the same 5-bp DR (5’-GTTTT-3’), and that they are inserted within another potential mobile element as judged by the external 34-bp inverted repeats (5’-GGGGGTGTAAGCCGGAACCCCAGAAAATTCCGTC-3’, gray triangles facing inwards) and accompanying direct repeats located upstream of the IRi and downstream of the IRt (Fig. 3). Our BLASTn search of the NCBI bacterial DNA database (as for October 9, 2020) using as query the pLD209 sequence (KF840720.1) found homology (99% nucleotide identity) between a composite fragment of 1,861 bp from this plasmid with a fragment extending 1,802 bp present in plasmid p3 from an environmental *Pseudomonas koreensis* strain, P19E3 (GenBank accession CP027480.1, positions 265,881 to 264,070). This region in *P. koreensis* p3 covered exactly an element bordered by identical 34-bp inverted repeats, and also similar hypothetical protein coding sequences, than those found in pLD209 after removing *in silico* the Tn*6335* insertion at the 5’-GTTTT-3’ direct repeat (122 bp from the IRi, from positions 891 to 1,012, and 1,739 bp from the IRt, from positions 8,651 to 10,389 in KF840720.1 (see also 13). These observations strongly suggest that a similar mobile external element was collected by a pLD209 ancestor, probably as the result of a *trans*-mediated transposition event, and subsequently targeted by a Tn*402*-like transposon thus generating the backbone of the adaptive module now observed in pLD209 and related plasmids (Fig. 3). These plasmids, which are amply disseminated among members of the *P. putida* group in clinical settings of Argentina, are endowed with the worrying potentiality to disseminate carbapenem and other antimicrobial resistance cassettes carried by Tn*402*-like integrons to co-existing pathogens including members of the *Enterobacteriaceae* family or *P. aeruginosa* as shown above.

**FIG 3.**
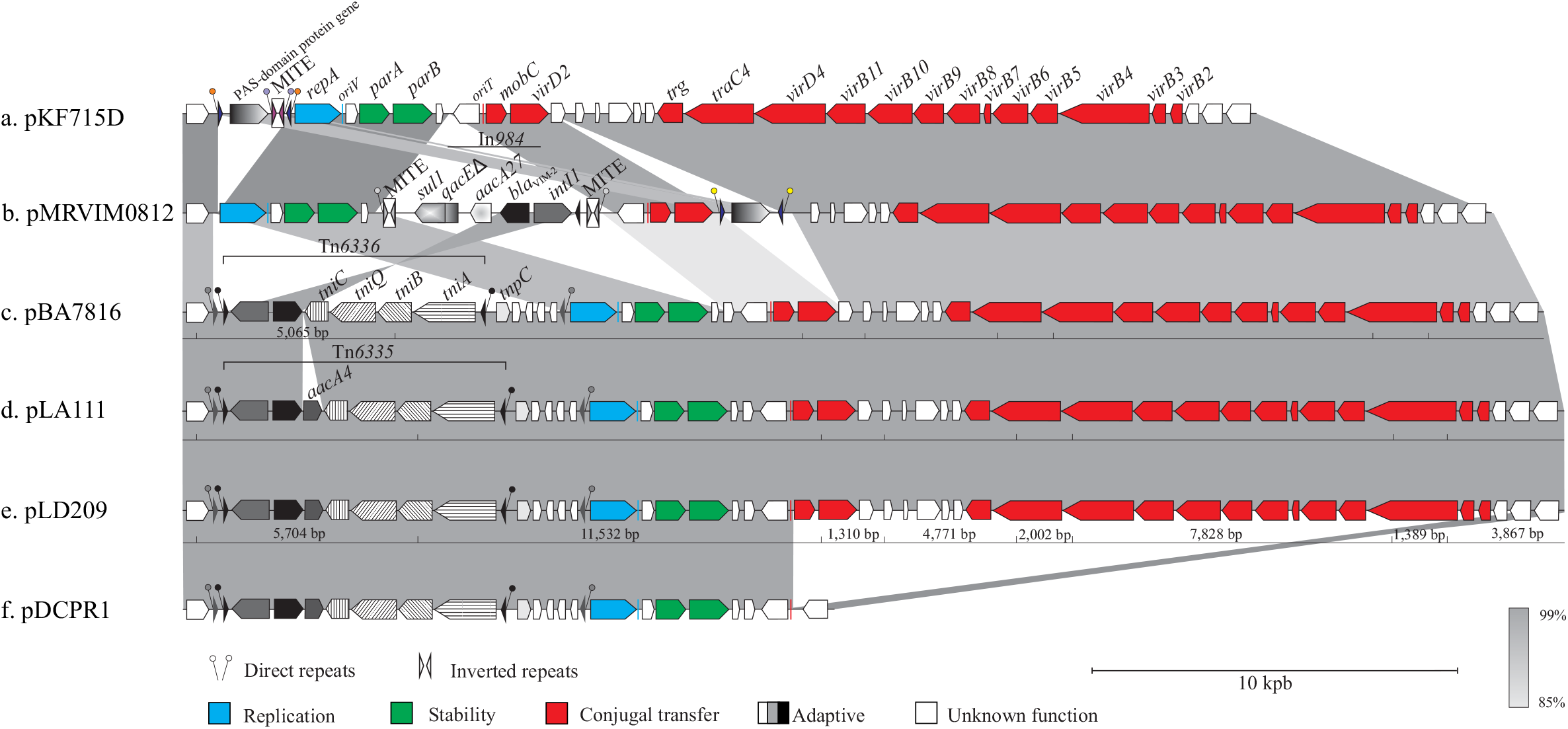
Comparative analysis of pLD209-type plasmids present in *P. putida* G strains. Linear representations of the structures of six circular plasmids including a) pKF715D (GenBank accession number AP015033.1), b) pMRVIM0812 (CP010893.1), c) pBA7816 (MN240297, this work), d) pLA111 (MT192131, this work), e) pLD209 (KF840720.1), f) pDCPR1 (KJ577613). The direction of transcription of the genes are indicated by arrows, and the different colors delineate the replication, stability and transfer modules (see the lower part of the figure). For the different structural components of Tn*6335* and Tn*6336*, including the adaptive gene cassettes present in the integrons they carry, see the legend to Fig. 2. The regions shaded in gray tones linking the different structures reflect percentages of nucleotide sequence identity ranging from 85% to 99% as detected in a BLASTn search, with the scale depicted at the lower right part of the figure. The positions of *Eco*RI restriction sites inferred from the corresponding DNA sequences are indicated below plasmids pBA7816, pLA111, and pLD209, with the fragment sizes predicted *in silico* shown below pLD209 only. In the case of pBA7816, the size of a differential EcoRI fragment when compared to the equivalent region in pLD209 (*i. e*., 5,065 versus 5,704 bp, respectively) is shown. A 5-bp 5’-GTTTT-3’ direct duplication (black circles) is present at the immediate outer borders of the 25-bp inverted repeats IRi and IRt (black triangles facing inwards accompanying the black circles) of both Tn*6335* and Tn*6336*. These Tn*402*-like elements are flanked by an external 34-bp inverted repeat (5’-GGGGGTGTAAGCCGGAACCCCAGAAAATTCCGTC-3’, gray triangles facing inwards), with one extreme (IRie) located immediately upstream of IRi and its complementary (IRte) located immediately upstream of the *repA* gene. These external IRe sequences are bordered by a 5’-TATTC-3’ direct repeat (gray circles accompanying the gray triangles). The single MITE element in pKF715D is located between nucleotide positions 20,221 to 20,482, and is flanked by a 5-bp direct repeat, 5’-AACTT-3’ (violet circles). The two MITE elements in pMRVIM0812 are located between positions 27,596-27,858 and 32,920-33,182, respectively, and the resulting composite transposon-like structure is flanked by the 5-bp direct repeat 5’-GATGA-3’ (light gray circles). The PAS-domain protein coding gene and adjacent MITE element in pKF715D are limited by a 50-bp inverted repeat, and flanked in turn by a 5-bp direct repeat, 5’-AGGAA-3’ (orange circles). In pMRVIM0812, the PAS-domain protein gene is also limited by a 50-bp inverted repeat flanked in turn by a 5-bp direct repeat, 5’-TGGAT-3’ (yellow circles).

Concerning the other seven *P. putida* G isolates analyzed, *i. e*., *P. asiatica* HP613, *P. putida* BA9115, *P. putida* BA7908, *P. monteilii* BA9713, *P. monteilii* HB157, *P. putida* G/I HP813, and *P. putida* G/II LA1008, our repeated attempts to detect the presence of plasmids by either conjugation or transformation assays were unsuccessful. This suggested a chromosomal, rather than a plasmid, location of the genetic elements harboring *bla*_VIM-2_ (Fig. 2) in these *P. putida* G isolates.

### Comparative analysis between pLD209 and related plasmids carried by nosocomial and environmental *Pseudomonas* species

Our BLASTn search using the pLD209 sequence as a query (see above) detected two plasmids, pKF715D and pMRVIM0812, showing high levels of nucleotide identity and structural organization with pLD209 including the replication, stability, and transfer modules (Fig. 3). Among them, pKF715D (Fig. 3a) was found in an environmental *P. putida* strain, KF715, obtained from contaminated soils near a biphenyl manufacturing plant in Japan (43, 44), and pMRVIM0812 (Fig 3b) was isolated from a clinical *Pseudomonas* sp. in the U.S.A.

At their adaptive modules, pMRVIM0812 contains a typical class 1 integron encompassing the *intI1* gene at the 5’-CS, *bla*_VIM-2_ and aminoglycoside 6’-acetyltransferase *aacA27* gene cassettes in the variable region, and the 3’-CS including *qacE*Δ-*sul1* genes (Fig. 3b). A similar class 1 integron, designated In*984*, was previously described in a clinical *Pseudomonas oleovorans* isolate, M13320 (42, GenBank accession number KJ668596). In pMRVIM0812 this class 1 integron is flanked by two copies of miniature inverted-repeat transposable elements (MITEs), the whole structure bordered in turn by a 5-bp (5’-GATGA-3’) DR (Fig. 3b). MITEs are non-autonomous mobile elements carrying inverted repeats, and their mobilization is promoted by transposases encoded by adjacent transposons or by ISs containing similar inverted repeats (45, 46). The assembly suggests that these MITE elements captured the integron in a composite transposon-like structure, which was subsequently mobilized to the present location in pMRVIM0812 by transposases provided in *trans*. Similar capturing and transposing events of other class 1 integrons by flanking MITE elements have been recognized as a mechanism for mobilizing antimicrobial resistance determinants (31, 47).

We noted that pMRVIM0812 and pKF715D, in contrast to other pLD209-type plasmids (Fig. 3c-e), carry each a PAS-domain protein gene (Fig. 3 a,b). In pMRVIM0812 the PAS-protein domain gene is bordered by a 50-bp inverted repeat, while in pKF715D is adjacent to a MITE element with the whole arrangement flanked in turn by a 50-bp inverted repeat. The inverted repeats in the above plasmids are flanked by 5-bp direct repeats in each case, *i. e*, 5’-AGGAA-3’ in pKF715D and 5’-TGGAT-3’ in pMRVIM0812 (Fig. 3 a,b). These analyses suggest that these PAS-domain coding sequences and associated elements reached their present plasmid locations assisted in *trans* by factors provided by mobile elements co-existing in the cells.

Finally, evidence exists that pLD209-related plasmids can additionally evolve not only by gaining or loosing individual resistance cassettes in the Tn*402* element, but also by losing significant parts of their backbones as exemplified by plasmid pDCPR1 (18,182 bp; Fig. 3f) isolated from clinical strains of both *P. aeruginosa* and *Serratia marcescens* in Argentina (48). pDCPR1 shares high structural and nucleotide similarity to pLD209 at the adaptive, replication, and stability modules, but lacks most genes involved in conjugal transfer (Fig. 3). Still, the retention of *oriT* sequences suggests a mobilization potentiality for pDCPR1 in the presence of a conjugative plasmid. The structural rearrangements found in pDCPR1 are most likely associated to lateral transfer, and reinforce the role of pLD209-related plasmids as efficient and plastic genetic platforms for the spreading of carbapenem and other antimicrobial resistance genes among nosocomial pathogens (13; 48).

The above observations disclose a wide dissemination of conjugative plasmids sharing similar backbones and structural organization among environmental and nosocomial members of the *Pseudomonas* genus. These plasmids are endowed with high adaptive significance, and have most likely collected different mobile elements and resistant determinants during transit through different bacterial hosts subjected to various selective conditions. Particular examples are pLD209 and related plasmids, which provide platforms endowed with lateral transfer ability to different class 1 integrons. These integrons apparently found their way to the plasmid structure either in the form of a Tn*402*-like transposon or other entities capable of transpose with the help of *trans*-acting factors. The plasticity inherent to these integron-bearing adaptive modules in terms of exchanging gene resistance cassettes, exemplified by the capturing of *bla*_VIM-2_ and other antimicrobial resistance genes (Fig. 3), has certainly contributed to the adaptation of *P. putida* G species to the challenges of the clinical setting. A worrying perspective thus emerges for the treatment of infections produced by MDR nosocomial pathogens such as *P. aeruginosa* or members of the *Enterobacteriaceae*, considering the ability of *P. putida* G species not only to serve as reservoirs but also disseminate these wide-host range resistance plasmids by horizontal transfer.

### *bla*_VIM-2_-containing Tn*402*-like transposons translocation to target sites in the genomes of *P. putida* G members

Database searching analysis and experimental evidences with plasmid model systems have identified a high selectivity of Tn*5053/*Tn*402* family members for targets clustered in, or close to, *res* regions upstream of *tnpR* genes of some members of the Tn*3* family, and the equivalent *par* regions associated to segregational mechanisms of particular plasmids (32, 33, 34, 35, 36, 37, 49). These studies also indicated that Tn*5053/*Tn*402* transposons generally insert on these target loci with the IRi boundary facing the resolvase gene, although in a very few number of cases the opposite orientation (with the IRt end more close to the recombinase gene) was also reported (26, 34, 35, 36). In addition, transposition events independent of the presence of a *res* locus have also been found, albeit at very low frequencies and involving random target selection and orientation (36, 50).

In six of the *P. putida* G strains analyzed here including *P. asiatica* (both Pa_A_ and Pa_B_ clones), *P. juntendi*, *P. putida* G/II, and *P. putida* G/V, the Tn*402*-like transposons were found in pLD209-type conjugative plasmids (Table 1). Our analysis above further indicated that these transposons are inserted in these plasmids into a defective element bordered by 34-bp inverted repeats, but no sequences resembling putative *res* or *par* target sites could be identified in the vicinity of the insertion site. This suggests either an unusual transposition event or, alternatively, substantial sequence rearrangements near the site of insertion as described in other cases (13, 33, 35, 51).

As noted above, Tn*6335* was found in the pLD209 plasmid in *P. asiatica* LD209, but apparently in another genomic location in isolate HP613 which is clonally related to LD209 (Table 1; Fig. 1 and Fig. S1). Different genomic locations for Tn*6335* were also found in *P. putida* G/II isolates HE1012 and LA1008 (Table 1; Fig. 1 and Fig. S1). Therefore, we decided to characterize in further detail the genomic context of Tn*6335* in both *P. asiatica* HP613 and *P. putida* G/II LA1008. For this purpose, the cloning of the genomic region near the IRi boundary of the transposon was attempted for both isolates, taking advantage of the presence of a nearby *bla*_VIM_-2 gene conferring ceftazidime resistance (Table S4) and the single EcoRI site within the *tniB* gene (Fig. 3). In line with this objective, we separately treated total genomic DNA from these isolates with EcoRI, ligated the digested products into EcoRI-digested pSU18 (52), transformed *E. coli* DH5α cells, and selected for colonies containing inserts that, besides the chloramphenicol resistance provided by pSU18, additionally conferred ceftazidime resistance (see Material and Methods for details). Different clones in each case were then subjected to plasmid purification and sequencing analysis of the inserts.

In the case of *P. asiatica* HP613, we recovered a 9,031-bp EcoRI insert (designated pSU18-HP613; Fig. S2A), showing the expected 5,307 bp region carrying *bla*_VIM-2_ and extending from the IRi of Tn*6335* to the EcoRI site located 406-bp of the 3’end of *tniB* (Fig. S2A). The remaining 3,725 bp falling outside the IRi boundary and extending to a nearby EcoRI site in the HP613 genome (Fig. S2A) exhibited 97.8% nucleotide identity to the indicated equivalent *Δres-tnpRA* region present in a complete Tn*501* element described in *P. aeruginosa* plasmid pVS1 (GenBank accession number Z00027.1, nucleotide positions 4,620 to 8,343; Fig. S2B). Tn*501* forms part of the Tn*21* subgroup of the Tn*3* transposon family (32, 33). All members of this subgroup show a *tnpRA* module composed of *tnpR* (serine recombinase) and *tnpA* (transposase) genes transcribed in the same direction, preceded by a *res* region composed of *resI, resII* and *resIII* subsites (32, 33, 53; 54; see also Fig. S2B). Our comparative sequence analysis indicated that in the *P. asiatica* HP613 genome the *resI* subsite of this Tn*501*-like element had in fact been impacted by Tn*6335*, with the IRi of the transposon facing the recombinase gene (Fig. S2B). A similar finding was observed by other authors in *P. aeruginosa* Pavimgi1, where the *res* site was also impacted between the *res*I and *res*II subsites (Fig. S2B) by another Tn*402*-like transposon (26).

A BLASTn search using this 4,442 bp genomic sequence as query identified almost identical stretches of around 3,790 bp (99 % nucleotide identity) in plasmid RPL11 of a *P. aeruginosa* isolate (GenBank accession AF313472) and in the chromosome of *P. putida* H8234 (GenBank accession CP005976.1); positions 3,173,668 to 3,178,100). This fragment includes a complete *tnpRA* module followed by a 38-bp IRr boundary (Fig. S2C) of a Tn*1403* transposon (Stokes et al. 2007). Tn*1403* is also included in the Tn*21* subgroup of the Tn*3* family (32). The remaining fragment towards the EcoRI site of around 650 bp also shows almost complete nucleotide identity to the equivalent region of the *P. putida* H8234 chromosome, and included 124 bp of the 3’ coding region of *sdhC* encoding the cytochrome b556 subunit of the succinate dehydrogenase (Fig. S2C). The complete *sdhC* gene (375 bp in length, GenBank accession AGN79094.1) is actually present in this locus in the *P. putida* H8234 chromosome, limited in turn at its 5’ region by another 38-bp inverted repeat characteristic of Tn*3* family transposons (not shown). This suggests that the whole element may in fact represent a novel Tn*1403*-like transposon carrying a *sdhC* catabolic gene. Similar genetic arrangements have in fact been described for other Tn*21* subgroups members of the Tn*3* family, although the catabolic genes they carry differ from the above (32).

Concerning the target region impacted by Tn*6335* in the *P. putida* G/II LA1008 genome, our comparative sequence analysis of the *res* regions of the Tn*1403*-like transposons mentioned above (Fig. S2D) indicated that Tn*6335* was inserted between the *resI* and *resII* subsites of the target element. As above, Tn*6335* was inserted with its IRi boundary facing the *tnpR* gene of the Tn*1403*-like target. The *tnpRA* transposition modules of the Tn*501*-like and Tn*1403*-like found above (Fig. S3) showed 86 % nucleotide identity between them, but differed in the length of the corresponding *tnpR* genes which were 561- and 618-bp, respectively. This situation has been described previously for Tn*21* subgroup members of the Tn*3* family (32, 53).

Finally, in the case of *P. monteilii* HB157 our analysis of the immediate genomic sequences in which the *bla*_VIM-2_-containing Tn*402*Δ(*tniABQ*) element (Fig. 2C) was located indicated that it was also inserted within the *res* region of another Tn*3* family transposon. Using an inverse PCR approach followed by cloning and sequencing of the obtained amplicons (see Materials and Methods for details), we could characterize a 732 bp fragment of the HB157 genome corresponding to the insertion site of Tn*402*Δ(*tniABQ*) in the immediate vicinity of its IRt border (Fig. S2E). Comparative sequence analysis indicated that this 732 bp fragment encompassed a complete *tnpR* resolvase gene (615 bp, 204 amino acids), followed by the first 14 bp of an aminoglycoside O-phosphotransferase (*aph*(*3*”)-Ib) gene (Fig. S2E). Moreover, a BLASTn search indicated that this fragment showed high identity with equivalent segments located in the genomes of different *Pseudomonas* species including (among others) the *P. aeruginosa* FDAARGOS_570 chromosome (Genbank CP033835.1, positions 3,824,646 to 3,825,377), a plasmid carried by the same strain (CP033834.1, positions 25,649 to 26380), the *P. mosselii* plasmid pMOS94 (MK671725.1, positions 21,091 to 21,822), the *P. putida* JBC17 chromosome (CP029693.1, positions 819327 to 823821), the *P. putida* 15420352 plasmid p420352-strA (MT074087.1, positions 126666 to 131160), as well as in the chromosomes and plasmids of other species of clinical and environmental relevance including *Klebsiella pneumoniae* PMK1 plasmid pPMK1-C; *Citrobacter freundii* RHBSTW-00444 plasmid pRHBSTW-00444_2; *Stenotrophomonas maltophilia* SM 866 chromosome; *Aeromonas caviae* WCW1-2 chromosome; *Aeromonas salmonicida* plasmid pRAS2, etc. In all of the above genomes this fragment forms part of a transposon found in *Aeromonas salmonicida* designated Tn*5393*c (55), which is essentially identical to the originally-described Tn*5393* found in *Erwinia amylovora* (56), an ubiquitously distributed member of the Tn*3* family carrying streptomycin resistance genes (32). In these Tn*5393*-like transposons the *tnpR* gene is separated from an oppositely-oriented *tnpA* gene (55) by a 125 bp intergenic region in which the three *res* subsites recognized by the recombinase (32, 57) could be inferred (Fig. S2F). Efficient transposition of Tn*5053*/Tn*402* members generally depend on externally coded accessory functions, namely a *res* site served by a cognate resolvase, but each interaction system may target a different *res* subregion (or even lie outside this region) (36). In this context, our analysis indicated that Tn*402*Δ(*tniABQ*) was inserted into the intergenic region between the *resII* and *resIII* subsites of a Tn*5393*-like element located in the *P. monteilii* HB157 genome (Figs. S2, E and F). Moreover, the IRt boundary of Tn*402*Δ(*tniABQ*) was found facing the *tnpR* resolvase gene of its Tn*5393*-like target (Figs. S2E and S3C), an insertion orientation different to the most frequently found for Tn*5053*/Tn*402* members including Tn*6335* on its Tn*21* targets described here (Fig. S2). This insertion orientation with the IRt closer to the *tnpR* of their targets has however been reported in other few cases for other members of the Tn*5053*/Tn*402* family (35, 36).

It has been suggested that the predisposition of Tn*5053*/Tn*402* family members for *res* sites provides access to alternative/more efficient vehicles of dissemination within different bacterial species sharing similar environmental niches, including those composing the human microbiota (32, 34, 58). Compound elements between Tn*402*-like and Tn*21* transposons, and subsequent derivatives, have in fact been found widely distributed in the human microbiome (58). The results presented here indicated that the *res* regions of Tn*21* transposons present in the genomes of different *P. putida* G species have been the preferential targets of Tn*402*-like elements carrying *bla*_VIM-2_ such as Tn*6335*, which most likely arrived to these cells as passengers of pLD209-related conjugative plasmids. Moreover, they also show that other members of the Tn*3* family such as Tn*5393*, of ubiquitous distribution among different gammaproteobacteria families, can also accommodate Tn*402*-like elements. It follows that *P. putida* G members can provide many elements to host Tn*402*-like integrons carrying antimicrobial resistance cassettes. Carbapenem therapy appears as a main force behind the selection of *P. putida* G clonal lineages in which transposition events relocated the incoming *bla*_VIM-2_-containing Tn*402*-like elements from plasmids to other preferred genomic locations such as some pre-existing members of the Tn*3* family, and also of further rearrangements occurring in the newly-generated hybrid structures. These chimeras combining Tn*402*-like and selected Tn*3* family elements could certainly play important roles in the dissemination of *bla*_VIM-2_ genes to pathogenic species of *Pseudomonas* and other bacteria causing infections in humans, food animals, and livestock (24, 26, 58, 59).

## DISCUSSION

We characterized here in detail the genetic platforms harboring *bla*_VIM-2_ in a set of carbapenem-resistant *P. putida* G clinical isolates obtained from different hospitals in Argentina, which were collected along an extended time period. Our study revealed notable taxonomic and genetic features among these isolates, which belong to a group better known to be composed of environmental organisms rather than nosocomial pathogens. The carbapenem resistant phenotype of our local collection of *P. putida* G isolates could be generally ascribed to the carriage of *bla*_VIM-2_ metallo-β-lactamase genes within Tn*402*-like integrons. The role of class 1 integrons in the dissemination of antimicrobial resistance among bacterial communities is well recognized (37), but the genetic elements that are exchanged between nosocomial and/or environmental bacteria and underlying mechanisms of dissemination still remain matters of debate and speculation. The detailed characterization conducted here of the genetic contexts in which *bla*_VIM-2_ was located in our *P. putida* G isolates thus helps our understanding of the events contributing to the spread of carbapenem resistance to pathogenic species in the nosocomial setting, and sheds light on the role of this bacterial group as an active environmental reservoir of antimicrobial resistance platforms.

The taxonomic characterization of the *P. putida* G isolates conducted first allowed us a better definition of the different species involved in this study. We were able to distinguish at the 13 *P. putida* G clinical isolates the species level, identifying among them representatives of *P. asiatica*, *P. putida sensu stricto*, *P. monteillii*, *P. juntendi*, and other 3 species more recently described as forming part of this group (1, 4, 5). In particular, our analyses (Table 1 and Fig. 1) could confidentially assign 4 of them including BA7816, LD209, HB313 and HP613 to the recently described species *P. asiatica* (4). Also, our comparisons indicated that the proposed species *P. putida* G/IV (2) can actually be ascribed to *P. asiatica* (Table 1 and Fig. 1). Our results also emphasize the ability of different members of the *P. putida* G to adapt and survive in the nosocomial habitat.

The searching of genetic elements containing *bla*_VIM-2_ conducted next among these *P. putida* G isolates revealed that three Tn*402*-like class 1 integrons (Fig. 3) carry this carbapenem resistance gene. In general, class 1 integrons are not mobile elements by themselves, but two described here, In*41* and In*899* (Fig. 2A) were found embedded in complete Tn*402*-like transposons (Tn*6335* and Tn*6336*, respectively) carried by pLD209-related plasmids (Fig. 3), and thus potentially capable of both intra- and inter--cellular mobilization (Fig. 4). In this context, the finding of identical 5-bp DRs bounding these transposons (indicated by black circles, Fig. 3, c-f) provides evidence that the original Tn*402*-like transposon from which these elements derive reached their plasmid location by a transposition event (32, 33, 34, 35, 36, 37, 49).

**FIG 4.**
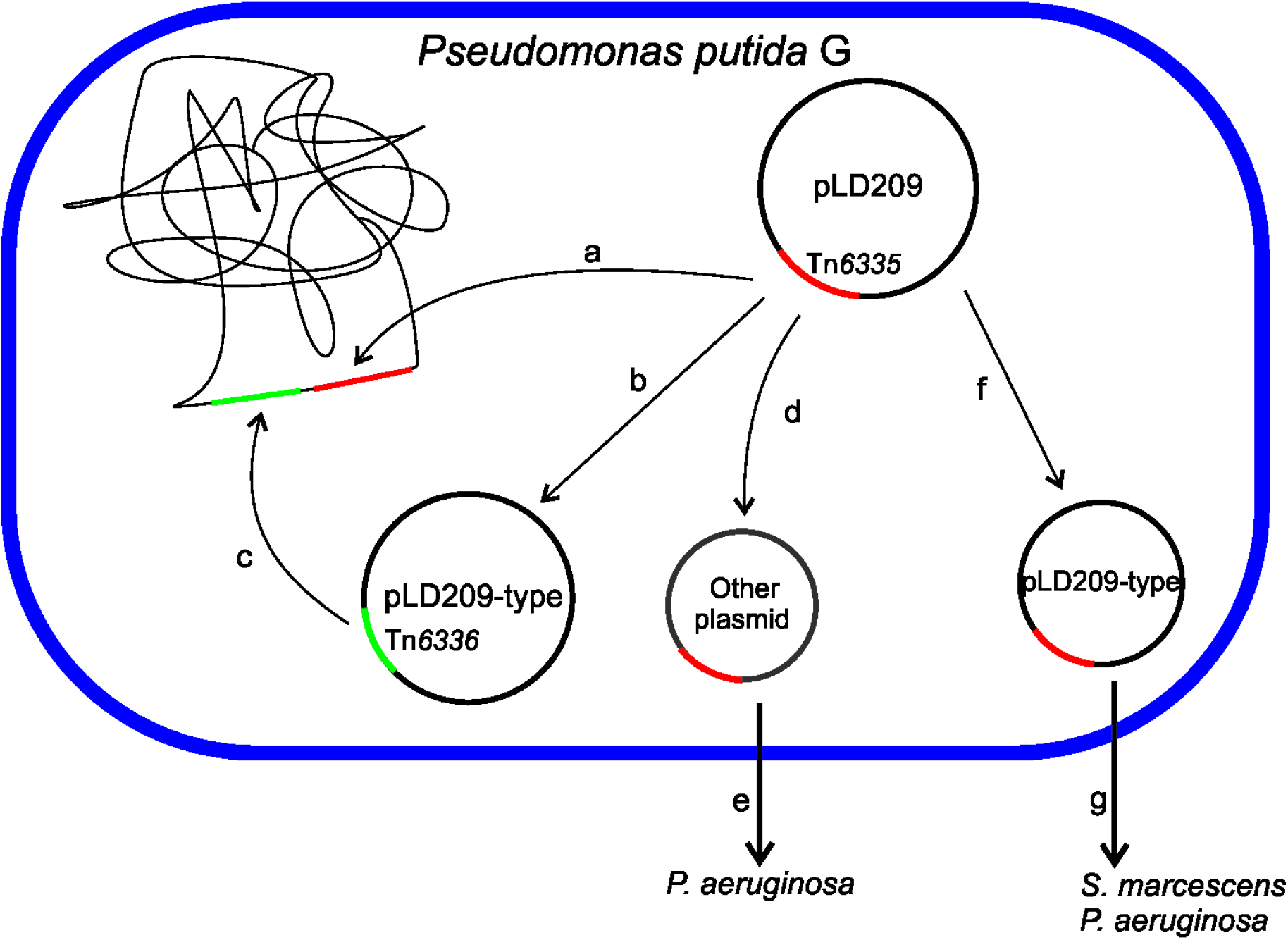
Routes of intra- and inter-genomic dissemination of *bla*_VIM-2_-containing genetic platforms among *P. putida* G members, and between this and other co-existing bacterial groups in the clinical setting. Tn*6335*, the prevalent Tn*402*-like transposon found in the *P. putida* G isolates studied here, was most probably acquired as a passenger of a conjugative pLD209-related plasmid. The plasmid could either persist in the new host, or fail to accompany the host replication rate and consequently be lost. In the latter case, carbapenem pressure could select bacterial clones in which the *bla*_VIM-2_-containing Tn*402*-like element had transposed from the plasmid to pre-existing preferred locations located in the chromosome or in other plasmids such as the *res* sites of Tn*21* subgroup transposons, exemplified in this work by the cases of *P. asiatica* HP613 and *P. putida* G/II LA1008 (a, c). Lack of aminoglycoside pressure could lead to the selection of clones in which the *aacA4* gene cassette was lost from the Tn*402*-like integron, resulting in a pLD209-related plasmid now harboring a Tn*6336* element (b), exemplified here in the case of *P. asiatica* BA7816 (Table 1). As above, clones in which this element had transposed to other preferred sites on the genome could be selected by carbapenem pressure, a situation exemplified here for *P. putida* BA9115 (Table 1) (c). The transposition of *bla*_VIM-2_-containing Tn*402*-like elements to a co-habitant plasmid displaying an idiosyncratic host range (d) also carries the possibility of a further dissemination of these elements by horizontal transfer to pathogenic bacteria, exemplified here by the finding of such a plasmid in a local *P. aeruginosa* clinical isolate, PAE868 (e) (see Discussion for details). Finally, deletions on pLD209 may have resulted in the selection of related plasmids lacking self-transferability, exemplified by pDCPR1 (Fig. 3) (f). Since pDCPR1 still preserved the *oriT* region, the Tn*402*-like element could still be transferred by conjugation if appropriate mobilization functions are provided in *trans* (g), exemplified by the isolation of this plasmid from both *P. aeruginosa* and *S. marcescens* clinical strains (48).

Besides their acquisition as passengers of pLD209-related conjugative plasmids, the selection of transposition events from the incoming plasmid to pre-existing preferred locations in the host genome, such as the *res* sites of Tn*21* subgroup transposons, provides another mechanism of *bla*_VIM-2_-containing Tn*402*-like integron dissemination among *P. putida* G members (Fig. 4, a and c). The selection of clones that had lost resistance cassettes such as *aacA4*, seemingly represents another genetic event occurring among *P. putida* G strains (Fig. 4b), as exemplified by the case of *P. asiatica* BA7816 harboring only *bla*_VIM-2_ into the integron variable region carried by a pLD209-related plasmid (Fig. 3). Similar to Tn*6335* above, clones may have been selected in which transposition of Tn*6336* to other genomic sites preserved the carbapenem resistance phenotype in the eventuality of pBA7816 loss (Fig. 4c), as exemplified in the case of *P. putida* BA9115 (Table 1). Another possibility is represented by the selection of plasmids that, with the exception of the replication and adaptive modules, have lost substantial portions of the original structures (Fig. 4f). Such pLD209-derived plasmids have in fact been recovered from clinical strains of *S. marcescens* and *P. aeruginosa* in local hospitals (Fig. 3, 48), suggesting that members of the *P. putida* G group could have been also their sources or reservoirs (Fig. 4, f and g). All these observations reinforce the existence of both assisted intra- and inter-cellular mobilization events of *bla*_VIM-2_-containing Tn*402*-like integrons among *P. putida* G species, increasing the threat of carbapenem resistance dissemination to other pathogenic species co-existing in the clinical setting. In the above context, we recently isolated a local *P. aeruginosa* clinical strain, PAE868 (Table S1), harboring a plasmid (pPAE868) carrying a Tn*6335* element. This plasmid has the ability to replicate in different *Pseudomonas* species including *P. aeruginosa* PAO1 and in the carbapenem-susceptible *P. juntendi* HPC451 strain characterized by us (Fig. 1, Table S3, and data not shown), but not in *E. coli*. These observations suggest that Tn*6335* could have been acquired by a pPAE868 predecessor during their co-existence in a same cell (Fig. 4d), prior to the horizontal transfer of the modified plasmid to *P. aeruginosa* (Fig. 4e).

The above observations reinforced our previous notion that self-mobilizable pLD209-type plasmids are capable of spreading *bla*_VIM-2_-containing Tn*402*-like integrons not only among a wide range of *P. putida* G species, but also to enterobacterial species (13, 48). In addition, the persistence and/or expansion of particular *P. putida* G clonal lineages such as Pa_A_ of *P. asiatica* (Table 1) certainly increases the possibilities of *bla*_VIM-2_ dissemination. In the latter context, we observed not only the presence of clonally-related *P. asiatica* isolates such as LD209 and HP613 in the same hospital at different dates (Table S1), but also an almost simultaneous presence of clonally-related isolates of this species in different hospitals such as HP613 and HB313 (Fig. S1 and Table S1).

In conclusion, our findings indicate that members of the *P. putida* G conform nowadays active parts of the nosocomial microbiota, representing an important reservoir of genetic determinants such as *bla*_VIM-2_ genes responsible of carbapenem resistance. Moreover, the findings here that particular Tn*402*-like class 1 integrons have been disseminating among *P. putida* G species in our clinical setting with the assistance of pLD209-type conjugative plasmids over a period of 9 years support the postulated ability of these mobile genetic elements to largely persist in the nosocomial habitat (60). Finally, the results presented here provide evidences supporting the intra- and intergenomic mobilization of Tn*402*-like integrons and derived composite transposons encompassing also Tn*3* family members among the components of this bacterial group, providing clues that may explain the vast dissemination of resistance genes among *Pseudomonas* and other pathogens.

## MATERIALS AND METHODS

### Bacterial isolates and antimicrobial susceptibility testing

A total of 13 carbapenem-resistant clinical isolates initially identified as *Pseudomonas putida* by the Vitek 2C System (bioMérieux, Marcy l’Etoile, France) were included in this study (Table 1). These isolates were collected from inpatients of different hospitals of Buenos Aires (B1-B3) or Rosario (R1-R5), Argentina, during the period 2006-2014 (Table S1). The isolates designated as BA were obtained from Instituto Malbrán, Buenos Aires. The susceptibilities to different antimicrobials including imipenem, meropenem, piperacillin-tazobactam, ceftazidime, cefepime, amikacin, gentamicin and ciprofloxacin of the different strains (Table S1) were evaluated usingns the Vitek 2C System (bioMérieux, Marcy l’Etoile, France). The interpretation of the MIC values shown in Table S1 was based on CLSI breakpoints recommendations (61).

### Assignments of *Pseudomonas putida* G clinical isolates to the species level and phylogenetic relatedness between strains and isolates

The assignment of each of the clinical isolates originally characterized as belonging to the *P. putida* group by phenotypic procedures (see above) to the species level was based on multilocus sequencing analysis (MLSA) and sequence comparisons following described procedures (1, 2). This approach is based on the percentage of nucleotide sequence identity between alignments of the concatenated sequences here employed (2,647 bp in total), corresponding to partial regions of the 16S rDNA (1,301 pb), *gyrB* (669 pb) and *rpoD* (677 bp) genes, and defined here as 97.5 % identity as the threshold value that separates species within the *P. putida* group. For this purpose, genomic DNA of each clinical isolate was purified using Wizard Genomic DNA Purification Kit (Promega, Madison, WI), and used as templates for PCR reactions aimed to amplify the desired fragments of the 16S rDNA, *gyrB* and *rpoD* genes used for concatenate construction (1, 2, 62; Table S2). The obtained amplicons were then sequenced at the Sequencing Facility of the University of Maine (Orono, ME, USA). The corresponding partial sequences of the mentioned genes from type strains including 21 species of *P. putida* G, *P. aeruginosa* ATCC 10145, and *P. oryzihabitans* ATCC 43272, as well as those corresponding to representative members of 6 newly *P. putida* G proposed species (2), were retrieved from the sequence data deposited on the NCBI database (Table S3). Alignments of concatenated genes were done using ClustalW with default parameters (https://www.genome.jp/tools-bin/clustalw). These alignments were also employed for the construction of a Maximum-Likelihood (ML) phylogenetic tree using MEGA7.0 (63). To determine the best-fit nucleotide substitution model, the tool included in MEGA7.0 was employed resulting in the use of the GTR+G+I substitution model, taking into account the Akaike information criterion (AIC). Only branches supported by bootstrap values higher than 60% (1,000 replicates) are shown in the depicted tree.

### Genomic relatedness among *P. putida* group isolates

The genomic relatedness among isolates assigned to the same species was evaluated by a random amplification PCR assay employing degenerate oligonucleotides (DO-PCR) (39).

### Detection of MβL by phenotypic and molecular methods

MβL-production was assayed by the EDTA-imipenem microbiological assay (EIM) and EDTA disk synergy test (EDS) (38). The presence of *bla*_VIM-like_, *bla*_IMP-like_, *bla*_SPM-1_ or *bla*_NDM-like_ genes was evaluated by PCR using specific primers (Table S2).

### Genetic environments of the *bla*_VIM-2_ genes in the *P. putida* G clinical isolates analyzed in this work

The association of *bla*_VIM-2_ genes with “unusual” class 1 integrons in the *P. putida* G clinical isolates studied here was detected by PCR using the primer pair 5’-CS (forward) and TniC-R2 (reverse) primers (Table S2) followed by sequencing analysis (17). The subsequent characterization of the structures of the Tn*402* structural elements in which the detected integrons were embedded (Fig. 2) was done by PCR-overlapping assays using genomic DNA from each isolate in each case and the appropriate pairs of primers (Table S2), followed by sequencing and database searching analyses as described in detail previously (17). Also, the left boundary sequences of the detected Tn*402* transposons from the IRi to the *bla*_VIM-2_ gene were determined by PCR using the primer combination IRHP/VIM-R; and the right boundary sequences from the *tniB* gene to the IRt by using the primer combination TniB-F/IRHP (Table S2). In the case of the Tn*402*-like element found in *P. monteilii* HB157, which lacks most of the *tni* module except for a complete *tniC* gene and 44 bp upstream of this gene (Fig. 2B), we completed the sequence of the whole element by an inverse PCR procedure (64). In short, 1 μg of genomic DNA from the corresponding isolates was digested by EcoRI (Promega, Madison, WI), the amplified DNA fragments were subjected to ethanol precipitation (65), resuspended in sterilized distilled water, and then ligated for 16 h at 4 °C with T4 DNA ligase (Promega). After a further purification step, PCR assays were performed in 25μl-reactions containing as template 0.1 μg of this circularized DNA, 0.5 μM of the primers VIM-Rf and IRHPr (Table S2), 200 μM of each dNTP, 2.0 mM MgSO4 and 1.0 U Platinum Taq DNA polymerase High Fidelity (Invitrogen, Carlsbad, CA). The cycling protocol involved 5 min denaturation at 94 °C, followed by 30 cycles of 30 s at 94 °C, 45 s at 53 °C, and 4 min at 68 °C, ending with a 10 min incubation at 68 °C. Several attempts using this procedure resulted in a discrete number of amplification bands ranging from around 2.5 to 4.5 kbp, probably as the result of secondary hybridization sites recognized by the primers employed. The amplification mixtures were thus subjected to ethanol precipitation and resuspension in sterilized distilled water as above, ligated to pGEM-T Easy (Promega), and transformed into *E. coli* DH5α cells by electroporation. After incubating the cells for 48 h at 37°C on LB agar plates supplemented with 100 μg/ml ampicillin, 40 μg/ml 5-bromo-4-chloro-3-indolyl-β-D-galactopyranoside (X-gal) and 54 μg/ml isopropyl β-D-1-thiogalactopyranoside (IPTG), plasmids were extracted from different colonies using the Wizard Plus SV Minipreps DNA Purification System, and analyzed by restriction mapping for the presence and size of inserts. The DNA sequences of selected inserts in the vicinity of the pGEM-T Easy cloning site were then determined to identify cloned fragments containing the desired sequences. We succeeded by this procedure in cloning an approximately 2.7 kbp DNA fragment, which initiated at the VIM-Rf primer (Table S2) and continued towards the *tniC* gene of the Tn*402* element (Fig. 2B). The sequence was completed by primer walking with the sequential use of tniC-F and IS6100-F primers (Table S2). This procedure allowed us to obtain the DNA sequence of a 2,758 bp fragment which not only covered the whole Tn*402* element (Fig. 2B), but also extended for an extra 732 bp into the *P. monteilii* HB157 genome in the immediate vicinity of its IRt border (Fig. S2E).

### Conjugation and transformation assays

Conjugation experiments were performed using the carbapenem-resistant *P. putida* G clinical isolates analyzed here as donors, and rifampicin-resistant cells of *E. coli* DH5α or *P. aeruginosa* PAO1 cells as recipients (17). Transconjugants carrying *bla*_VIM-2_-containing plasmids were selected on LB agar containing 20 μg/ml ampicillin and 150 μg/ml rifampicin in the former case, or 4 μg/ml ceftazidime and 150 μg/ml rifampicin in the latter. MIC values towards different antimicrobials were determined on the obtained *E. coli* DH5α transconjugants as described above. The subsequent self-transferability of the plasmids present in the *E. coli* DH5α transconjugants was further tested by agar mating studies employing as recipient the *E. coli* MC4100 strain harboring the chloramphenicol-resistance plasmid pACYC184 (66). Transconjugants were selected in these cases using LB agar plates containing 20 μg/ml ampicillin and 25 μg/ml chloramphenicol, and the loss of rifampicin resistance was confirmed in all cases.

In the cases of *P. putida* G isolates in which no *E. coli* transconjugants could be obtained by using the above procedures, plasmid DNA was isolated using the Wizard Plus SV Minipreps DNA Purification System and used to transform *E. coli* DH5α which had been made competent by chemical (CaCl_2_) procedures (65). Colonies were then selected on LB agar plates containing 20 μg/ml ampicillin, after an overnight incubation at 37 °C. *P. aeruginosa* PAO1 transformation was conducted by electroporation following described procedures, followed by selection of resistant colonies in LB agar containing 4 μg/ml ceftazidime (67).

The actual presence of the *bla*_VIM-2_ gene in the putative transconjugant or transformant cells obtained as described above was confirmed by PCR and sequencing analysis (Table S2). Plasmids from *E. coli* DH5α transconjugants were purified using the Wizard Plus SV Minipreps DNA Purification System, and further characterized by EcoRI digestion followed by agarose gel (0.7%) electrophoresis analysis of the obtained fragments (68). Selected plasmids were subjected to further sequencing analysis (see below).

### Plasmid sequencing and comparative sequence analyses

pLA111 and pBA7816 nucleotide sequences were determined on a 454 pyrosequencing platform (Roche Diagnostics Corporation) at the Instituto de Agrobiotecnología Rosario (INDEAR). The obtained reads were assembled *in silico* using as framework the structure previously determined for pLD209 (13). The circular structures of these plasmids were confirmed by PCR procedures, in which remaining gaps between the resulting contigs were closed using specifically designed primer pairs (Table S2). In the case of pLA111 we employed a virB10-F/virB8-R primer combination, and for pBA7816 we used mob-F/mob-R, 7816-F/VIM-R and TniA-F2/REPIR-T3 combinations (Table S2).

The DNA sequences of all the PCR amplicons and cloned inserts obtained in this work were done at the University of Maine DNA Sequencing Facility, Orono, USA.

The Rapid Annotation using Subsystem Technology standard operating procedures (RAST, http://rast.nmpdr.org/seedviewer.cgi) (69) and the National Center for Biotechnology Information database (NCBI, U.S. National Library of Medicine, Bethesda MD, USA) were used to annotate the open reading frames (ORFs). Searching for antimicrobial resistance determinants was done using ResFinder 2.1 (https://cge.cbs.dtu.dk/services/ResFinder/; 70). The detection of IS was done with ISFinder (71) (https://www-is.biotoul.fr/) and ISsaga (72). The sequences of pBA7816 and pLA111 were deposited at the GenBank nucleotide sequence database under the accession numbers MN240297 and MT192131, respectively.

### Target sites of Tn*6335* in the chromosome of selected *P. putida* G isolates

Genomic DNA was purified from *P. asiatica* HP613 and *P. putida* G/II LA1008 using the Wizard Genomic DNA Purification Kit, separately digested with EcoRI and ligated to EcoRI-digested pSU18, a *E. coli* cloning vector conferring chloramphenicol resistance (52). The ligation mixture was transformed into *E. coli* DH5α by electroporation, and transformants carrying inserts containing complete *bla*_VIM-2_ genes were selected on LB agar plates containing ceftazidime (4 μg/ml) and chloramphenicol (25 μg/ml) supplemented with 40 μg/ml X-gal and 54 μg/ml IPTG. After a 48 h incubation at 37 °C, plasmids were recovered from selected colonies as described above, and the DNA sequences of the cloned EcoRI fragments were determined employing first a primer hybridizing in the multiple cloning site of pSU18 (pSU18-F, Table S2), followed by primer walking using the sequence information obtained in each case.

## ACKNOWLEDGMENTS

We are grateful to the personnel of the Bacteriology Service, Hospital Provincial, Rosario, Argentina for kindly providing *P. putida* G clinical isolates used in this work. P.M. and A.L. are Researchers of the National University of Rosario. A.M.V. and D.F. are Careers Researchers of CONICET. M.B. is Fellow of CONICET. F.P. and A.C. are Researchers of the Malbrán Institute, Buenos Aires. This work was supported by grants from the Agencia Nacional de Promoción Científica y Tecnológica (ANPCyT; PICT 2012-0680 to A.L., and PICT 2015-1072 to A.M.V.); Consejo Nacional de Investigaciones Científicas y Técnicas (CONICET); Secretaría de Ciencia, Tecnología e Innovación, Provincia de Santa Fe, and Secretaría de Salud Pública, Municipalidad de Rosario.

---

**FIG S1. Identification of different clonal lineages within particular species of the *P. putida* group assigned in this work by random PCR assays.** Random PCR assays with degenerate oligonucleotide primers were conducted as described in Materials and Methods. Lanes 1 and 2: Pp_A_ and Pp_B_ clonal lineages identified in the two indicated *P. putida sensu stricto* isolates; lanes 3 and 4: Pm_A_ and Pm_B_ clonal lineages identified in the indicated *P. monteilii* isolates; lanes 5 and 6: PpG/IIA and PpG/IIB clonal lineages identified in the indicated *P. putida* G/II isolates; lanes 7-10: two different clonal lineages, Pa_B_ (lane 7) and Pa_A_ (lanes 8-10) identified in the indicated *P. asiatica* isolates.

**FIG S2. Schematic representations of Tn*402*-like transposon insertions detected in this work into the *res* sites of Tn*3* family members in the genomes of *P. asiatica* HP613, *P. putida* G/II LA1008, and *P. monteilii* HB157. A.** Structural features of the 9,031 bp-*Eco*RI fragment containing a partial fragment of Tn*6335* carrying *bla*_VIM-2_ cloned from *P. asiatica* HP613. The fragment encompasses the 5,307 bp region of Tn*6335* spanning from the IRi to the EcoRI site present within the *tniB* gene (see Fig. 3), plus a 3,725 bp-fragment outside the IRi boundary to a nearby EcoRI site located in the HP613 genome. The latter fragment shows 97.8% nucleotide identity to the indicated *tnpA-tnpR* module and partial *res* site (□*res*), located in a complete Tn*501* element described in *P. aeruginosa* plasmid pVS1 (GenBank accession Z00027.1) (see legend to Fig. S3 for sequence details and also the main text). **B.** Alignments of the ~200 bp homologous *res* regions (including the remaining *resI* as well as *res*II and *res*III subsites) located upstream of *tnpR* recombinase genes of Tn*501*-like elements impacted by Tn*6335* in HP613 (pSU18-HP613, this work) and by another Tn*402*-like transposon in *P. aeruginosa* Pavimgi1 (KJ463833.1, 26, included for comparison purposes). The equivalent complete *res* region of a Tn*501* element present in plasmid pVS1 (see the main text) is also shown at the bottom, for a better appreciation of the *res* subsites locations and the precise sites of insertion (black arrowheads) of the Tn*402* transposons in the above cases. Conserved nucleotides between the three *res* regions are indicated by asterisks below the sequences, and the different *res* subsites are boxed. All sequences are shown in the 5’ to 3’ direction corresponding to the *tnpR* coding strand, and the position of the *tnpR* translation initiation codons is also indicated above the alignments. **C.** Structural features of the 9,749 bp-*Eco*RI fragment containing a partial fragment of Tn*6335* carrying *bla*_VIM-2_ cloned from *P. putida* G/II LA1008. The fragment encompasses the Tn*6335* region from the IRi to the *tniB* EcoRI site as above, plus a 4,442 bp fragment outside the IRi boundary to a nearby EcoRI site located in the LA1008 genome. The latter fragment shows 99% nucleotide identity to the indicated region of a Tn*1403*-like transposon and neighbouring regions (see Fig. S3 for sequence details and also the main text) outside the IRr in the *P. putida* H8234 chromosome (GenBank CP005976.1; positions 3,173,668 to 3,178,100; 73). **D.** Alignments of the ~120 bp *res* regions located upstream of the *tnpR* genes of the Tn*1403*-like elements located in the genomes of LA1008 (pSU18-LA1008, this work), *P. putida* H8234, and *P. aeruginosa* plasmid RPL11 (AF313472). The insertion sites of the Tn*402*-like transposons in LA1008 and RPL11 are indicated by black arrowheads. In plasmid RPL11, the *resI* subsite was reconstructed by fusing the 5’-AACTG-3’ DR sequence (labeled in dark gray) corresponding to the site where a Tn*402*-like element was inserted (AF313472). As above, all sequences are shown in the 5’ to 3’ direction corresponding to the *tnpR* coding strand, with the ATG translation initiation codon indicated above the sequences. Conserved nucleotides between sequences are also indicated by asterisks below the sequences. **E.** Structural features of the 5,971 bp-region containing the Tn*402*Δ(*tniABQ*) element inserted into the *res* region of a Tn*5393*-like element located in the *P. monteilii* HB157 genome. The complete sequence was derived from the combined data of PCR assays, inverse PCR, and cloning procedures (see Materials and Methods for details). The fragment encompasses the 5,239 bp Tn*402*Δ(*tniABQ*) element spanning from the IRi to the IRt boundaries (Fig. 2B), plus a 732 bp-fragment of the HB157 genome located outside the IRt that included the *resIII* subsite, the complete *tnpR* gene, and the first 14 bp of an *aph*(*3*”)-Ib gene. The latter fragment shows complete nucleotide identity to the equivalent region of a Tn*5393*c element described in *Aeromonas salmonicida* subsp. *salmonicida* plasmid pRAS2 (AF262622.1; 55). **F.** Alignments of the 125 bp *tnpR-tnpA res* intergenic region containing the *resI, resII* and *resIII* subsites of the *A. salmonicida* pRAS2 Tn*5393c* transposon with the remnant equivalent region impacted by Tn*402*Δ(*tniABQ*) in the HB157 genome. The precise site of insertion of Tn*402*Δ(*tniABQ*), 38 bp upstream of the *tnpR* gene of a Tn*5393*-like element located in this genome, is shown by a black arrowhead. The Tn*402* element is inserted immediately upstream of the *resIII* subsite of the Tn*5393*-like element, with the IRt facing the *tnpR* gene. Conserved nucleotides are indicated by asterisks below the sequences, and the different *res* subsites are boxed. All sequences are shown in the 5’ to 3’ direction corresponding to the *tnpR* coding strand, and the *tnpR* translation initiation codons are indicated above the alignments. The *res* subsites of the Tn*501*-like and Tn*1403*-like transposons were delineated as in ref. 54; and those of the Tn*5393*-like elements using the Tn*3* subsites following ref. 57.

**FIG S3. Nucleotide sequences and structural features of the target sequences corresponding to Tn*3* family transposons in which the Tn*402*-like transposons described in this work were inserted. A.** DNA sequence the EcoRI fragment cloned into pSU18 (pSU18-HP613) corresponding to the *tnpRA* region and remaining *res* region of the Tn*501*-like element impacted by Tn*6335* in the *P. asiatica* HP613 genome. **B.** Same, for the Tn*1403*-like element present in the *P. putida* G/II LA1008 genome (pSU18-LA1008). **C.** DNA sequence of the 732 bp-fragment of the *P. monteilii* HB157 genome in the immediate vicinity of the IRt of the Tn*402*Δ(*tniABQ*) element. All sequences are shown in the 5’ to 3’ direction corresponding to the direction of transcription of the *tnpR* recombinase genes.

